# Identifying and ranking potential driver genes of Alzheimer’s Disease using multi-view evidence aggregation

**DOI:** 10.1101/534305

**Authors:** Sumit Mukherjee, Thanneer Perumal, Kenneth Daily, Solveig Sieberts, Larsson Omberg, Christoph Preuss, Gregory Carter, Lara Mangravite, Benjamin Logsdon

## Abstract

**Motivation:** Late onset Alzheimers disease (LOAD) is currently a disease with no known effective treatment options. To address this, there have been a recent surge in the generation of multi-modality data (Hodes and Buckholtz, 2016; Mueller *et al*., 2005) to understand the biology of the disease and potential drivers that causally regulate it. However, most analytic studies using these data-sets focus on uni-modal analysis of the data. Here we propose a data-driven approach to integrate multiple data types and analytic outcomes to aggregate evidences to support the hypothesis that a gene is a genetic driver of the disease. The main algorithmic contributions of our paper are: i) A general machine learning framework to learn the key characteristics of a few known driver genes from multiple feature-sets and identifying other potential driver genes which have similar feature representations, and ii) A flexible ranking scheme with the ability to integrate external validation in the form of Genome Wide Association Study (GWAS) summary statistics. While we currently focus on demonstrating the effectiveness of the approach using different analytic outcomes from RNA-Seq studies, this method is easily generalizable to other data modalities and analysis types.

**Results:** We demonstrate the utility of our machine learning algorithm on two benchmark multi-view datasets by significantly outperforming the baseline approaches in predicting missing labels. We then use the algorithm to predict and rank potential drivers of Alzheimers. We show that our ranked genes show a significant enrichment for SNPs associated with Alzheimers, and are enriched in pathways that have been previously associated with the disease.

**Availability:** Source code and link to all feature sets is availabile at *https://github.com/Sage-Bionetworks/EvidenceAggregatedDriverRanking*.

## 1 INTRODUCTION

Late onset Alzheimers disease (LOAD) is a debilitating illness with no known disease modifying treatment (Alzheimers, 2015; Frozza *et al*., 2018). Identification new genetic drivers of LOAD will be key to the development of effective disease modifying therapeutics. To prioritize experimental evaluation of LOAD drivers, we present a data driven approach to rank genes based on the probability that they drive LOAD using transcriptional (RNA-seq) data collected from postmortem brain tissue in patient cohorts.

While there exists some prior work on driver gene ranking (Mukherjee *et al*., 2018; Hou and Ma, 2014; Liu *et al*., 2015; Grechkin *et al*., 2016; Zhang *et al*., 2013), these approaches have several limitations that make them unsuitable for all feature types. Many of these approaches work only with somatic mutation data from patients tumor samples, ranking genes by comparing the mutation rates of somatic variants in patients for different genes to an appropriate null model to identify cancer driver genes (Tian *et al*., 2014). While some other approaches use ensemble approaches to rank c.f.r genes using predictions from other tools that use genomic data (Liu *et al*., 2015). Unfortunately these approaches are highly specialized to the type of data and cannot be easily generalized to a broader class of feature sets. While there exist approaches such as DawnRank (Hou and Ma, 2014) which utilize RNA-Seq data in addition to genomic data for each patient, these too have strong modeling assumptions leading to lack of generalizability. Furthermore, most of these previous approaches are designed for detecting driver genes that are driven by somatic mutation events aside from the Key Driver analysis of (Zhang and Zhu, 2013). Alternatively, we are interested in identifying signatures of driverness from somatic tissue that are indicative of germline risk for LOAD. Here we propose a highly generalizable machine learning approach to learn signatures of germline genetic risk within summaries of tranascriptomic expression of somatic post-mortem brain tissue driver ranking and demonstrate it’s effectiveness on RNA-Seq derived featuresets.

Our driver ranking approach serves as an evidence aggregation framework, and currently uses differential expression, undirected gene networks inferred with an ensemble coexpression network inference method and co-expression module summaries (Logsdon *et al*., 2019) generated using transcriptional data collected from postmortem brain tissue across three studies (ROSMAP, Mayo RNAseq, MSBB) in AMP-AD. We assume that each analytic summary (while originating from the same RNA-seq data-sets) contains independently predictive information that can be used to identify genes with a burden of germline AD risk variants. We process these independent analytic summaries into the following feature sets (see Table 1) to be used for machine learning: 1) genes that are differentially expressed between AD cases and controls in specific brain regions, 2) global undirected network topological features for specific brain regions, and 3) module specific network topological features for 42 tissue specific co-expression modules.

**Table 1.** Description of various feature sets used for multi-view evidence aggregation.

| Feature Set | SynapseID | No. features | Type | Description |
| --- | --- | --- | --- | --- |
| Differential Expression | syn18097426 | 250 | Binary | Membership based on differential expression in different brain regions and patient sub-groups (such as males/females) |
| Global-Network | syn18097427 | 42 | Numeric | Features derived from graph structure in different brain regions |
| Module-Network | syn18097424 | 66 | Numeric | Features derived from graph structure in important co-expression modules from different brain regions |

Here we divide the task of ranking potential driver genes into two sub-tasks: i) training machine learning models to identify probabilities of genes being driver genes using each feature set, ii) aggregation of predictions of models for each feature set along with independent GWAS statistics to rank potential driver genes (Figure 1). The primary goal of the first task is to learn the unique characteristics of 27 previously known drivers of AD identified from published LOAD GWAS studies (Lambert *et al*., 2013; Kunkle *et al*., 2018) and use it to identify potential novel drivers of the disease. These AD drivers were defined as loci that were genome-wide significant in one study (P¡5×10-8), with significant replication p-value (P¡0.05) in a second study. The technical challenges associated with the first task include finding an appropriate approach to identify the driver probabilities and finding a way to learn from sparsely labeled data (only 27 genes have labels, while others may or may not be driver genes). To tackle this, here we propose a novel multi-view classification (Xu et al., 2013) approach, which includes iterative update of labels to infer additional candidate driver genes. For the latter task the primary challenge is to define an appropriate scoring system to rank genes. Here we propose a flexible scoring system that not only utilizes model predictions for each feature set but also independent LOAD Genome Wide Association Study (GWAS) statistics.

**Fig. 1.**
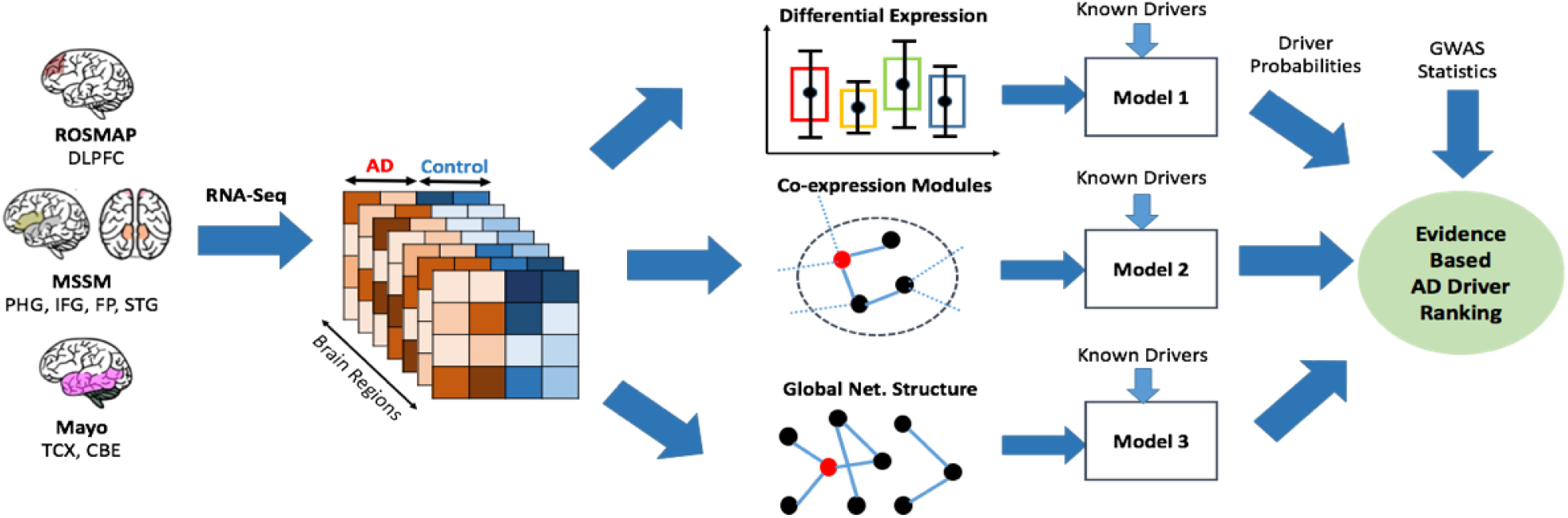
RNA-Seq data for AD patients and controls were derived for 7 different brain regions from 3 centers. Differential expression, Co-expression module, and Global network features were derived from all brain regions. Each feature and known drivers were used to build predictive models for driver genes. These driver probabilities and GWAS statistics were used for an evidence-based driver ranking.

We demonstrate our multi-view classification algorithm achieves substantially higher performance compared to models trained for individual feature sets on standardized multi-view datasets. We then demonstrate that similar performance benefits hold when applied to LOAD post-mortem brain tissue RNA-seq using qualitative metrics. We observe that global network topological features from inferred sparse coexpression networks - such as node degree - are predictive of LOAD driver genes as identified in GWAS, and more so than differential expression features. Finally, we show that our ranking methodology identifies several previously known LOAD loci implicated in other studies (Mukherjee *et al*., 2017; Ki *et al*., 2002; Kiyota *et al*., 2015; Jonsson *et al*., 2013) as well potentially new LOAD risk loci. These findings may lead to new mechanistic hypotheses regarding the genetic drivers of LOAD. Furthermore, a Gene Ontology (Chen *et al*., 2013) pathway analysis of the highly ranked predicted driver genes identifies multiple pathways previously implicated in LOAD disease etiology.

## 2 METHODS

### 2.1 Study description

Briefly, all feature sets are derived from analyses of RNA-seq data on 2114 samples from 1100 patients from seven distinct brain regions (Temporal Cortex, Cerebellum, Frontal Pole, Inferior Frontal Gyrus, Superior Temporal Gyrus, Parahippocampal Gyrus, Dorsolateral prefrontal cortex) and three studies - the Mount Sinai Brain Bank study (Wang *et al*., 2018), the Mayo RNAseq study (Allen *et al*., 2016), and the ROSMAP study (A Bennett *et al*., 2012). A full description of the data and the RNA-seq processing pipeline that was used to generate analytic outputs is described in (Logsdon *et al*., 2019).

### 2.2 Deriving usable features for meta-analysis

Features were inferred from specific statistical analyses that were run on RNA-seq data-sets within each of the seven tissue types. These analyses included set membership features from differential expression analysis (e.g. test of changes in mean expression between AD cases/controls and sub-groups such as males and females), global network features from a sparse ensemble coexpression network inference method described in further detail in (Logsdon *et al*., 2019), and network topological features for communities of genes identified from the networks described in the same paper. The sparse network inference approach applies 17 distinct coexpression network inference algorithms to data derived from each tissue type, and averqages across them to determine an ensemble sparse representation of coexpression relationships. In all network type features we extract standard network topological characteristics such degree, authority score, betweeness centraility, pagerank, and closeness.

### 2.3 Iterative multi-view classification for driver prediction

Here we pose the driver gene prediction as a binary classification problem using corrupted labels (Frénay and Verleysen, 2014). Formally, given a feature vector *X_i_* ∈ **R**^*d*^ for a gene denoted by the index *i*, we wish to predict a class label from {0,1} where 1 would indicate that the gene is a driver gene and 0 if it’s not. Additionally, we also desire to predict the conditional probability for of a gene being a driver, given the feature information i.e. ℙ(*Ŷ_i_* = 1|*X* = *X_i_*). This problem is solved by a broad class of binary classificaiton problems such as Logistic Regression, Support Vector Machines (SVM) etc. in the presence of a training dataset with input features and output class labels. However, here we are only provided a list of a small subset of drivers (from existing literature) whereas all other genes may or may not be a driver. Mathematically this is akin to learning from noisy labels *Ỹ* instead of the actual labels *Y* where ℙ(*Y* = 1|Ỹ = 1) = 1 but ℙ(*Y* = 0|Ỹ = 0) ≠ 1. While there are many general strategies for learning from noisy labels such as removing bad data points, active learning etc. (Frénay and Verleysen, 2014), they generally don’t account for this specific type of label noise or make assumptions about rates of mis-labeling in each class. Hence, here we focus on a simple existing approach for such problems (Iterative Classification) and propose a variant of it utilizing the fact that we have features from multiple views for the same genes.

#### 2.3.1 Iterative Classifier (IC)

Iterative classification is a simple approach where the general idea is to update the labels samples where *Ỹ* = 1 to that of the predicted class *Ŷ* after each iteration of model training (Liu *et al*., 2003). This can be written in algorithmic terms as in Algorithm 1. While this algorithm is general and can be used for different classifiers, here we demonstrate it on a L2-penalized Logistic Regression. Here, *ll* denotes the Maximum Likelihood loss for Logistic Regression and *thresh* is a constant in [0,1], typically chosen to be greater than 0.5. The higher the threshold, the more conservative the iterative updates are, acting as a trade-off between specificity and sensitivity.

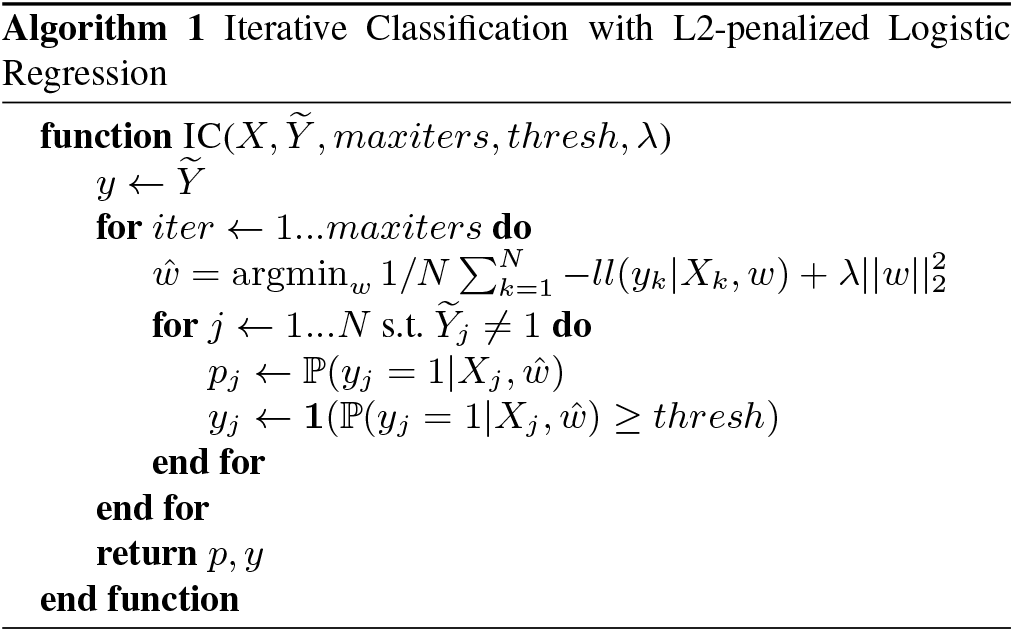

In the presence of data from multiple views from the same samples 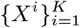, the algorithm is run for each view separately and an average of the predicted probabilities of all models is considered while evaluating the final multi-view predictions (we shall refer to this as ‘consensus’ for short in later text and figures).

#### 2.3.2 Iterative Classifier with Co-training (ICCT)

While the previous algorithm solves the problem of noisy labels and integrates information from multiple views, it does so by training models for each individual view independently. However, as seen in Figure 1, the features for different views are generated from the same underlying source i.e. the RNA-Seq data from brain samples of patients and controls. Hence, the different views can be seen as functional transformations of the same underlying data, corrupted with different noise sources and should encode the same classification information.

In the case of original multi-view classification problems, it is common to enforce view similarity which requires predictions made by different views to be similar to each other, through co-training or co-regularization (Xu *et al*., 2013). Here, the problem is more difficult to the noise in the labels. Hence, we develop a method which integrates the iterative updating scheme developed previously with co-training. Formally, we pose the problem of iteratively learning labels with co-training as the following optimization problem:

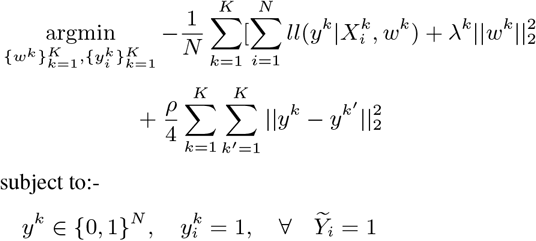

It can be seen that this is a mixed-integer optimization problem, which is a particularly hard class of optimization problems to solve. However, for fixed 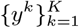, the optimization problem is convex in 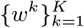 and is simply logistic regression for the different views. Hence, a potential solution to the optimization problem is via alternative minimization on 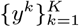 and 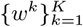 starting with 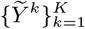. Unfortunately, the problem of optimizing over 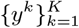 is a constrained binary quadratic programming problem, which does not have exact solutions or efficient exact solvers (Kochenberger *et al*., 2014). However, upon relaxing the binary constraint to a linear constraint ({0,1} → [0,1]), the optimization problem becomes a tractable convex optimization problem:

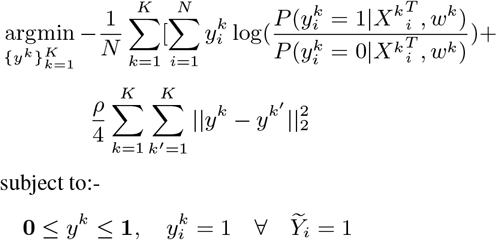

Here we note that 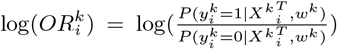. We note that this optimization problem is independent in each *i* and can be solved independently. Next we demonstrate that the previously posed linear relaxation which can be solved using the co-ordinate descent methodology using a closed form update rule for each 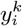.

##### Claim 1

A co-ordinate descent strategy leads to an optimal solution to the previously stated optimization problem.

**Proof**: It is sufficient to show that the optimization problem is convex. Since the inequality constraints are linear in 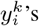, to demonstrate convexity of the optimization problem, we simply need to demonstrate that the cost function is convex. This can be shown by re-parameterizing the problem for the *i*th variable in terms of a new variable 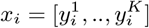.

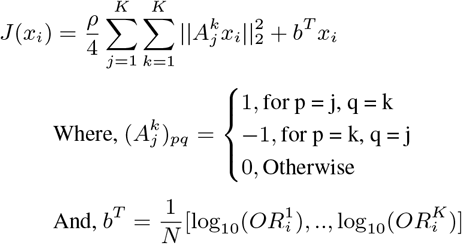

Next we calculate the second derivative of *J*(*x_i_*):

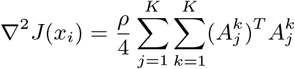

We see that, this is a sum of positive semi-definite matrices, ∇^2^*J*(*x_i_*) ≿ 0 for all *x_i_*, which is a sufficient condition for convexity (Q.E.D.).

##### Claim 2

The previously stated optimization problem has a closed form co-ordinate descent rule given by:

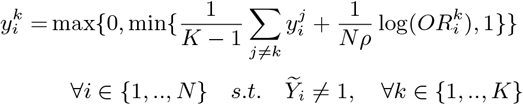

**Proof**: The loss function for each 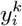 can be written as:

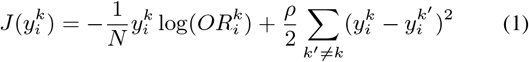

It is easy to see that this is a parabola of the form *y* = *a*(*x* - *b*)^2^ + *c*. For a parabola of this form, the minima (if *a* > 0) or maxima (if *a* < 0) occurs at *x* = *b*. For our cost function, we see that 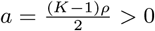 and 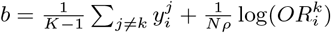. Hence, 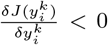 if 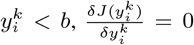 if 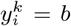 and 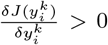 if 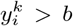. We now look at three possible locations of 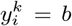 with respect to the interval 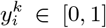 and the constrained minima in each case:

*Case I (b* ∈ [0,1]): Here the constrained minima is the same as the global minima.
*Case II (b* < 0): Here, 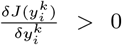 in [0, 1]. Hence, the constrained minima occurs at 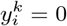.
*Case III (b* > 0): Here, 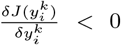 in [0, 1]. Hence, the constrained minima occurs at 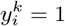.

Now, compiling the closed form solutions in the three cases, we can re-write the co-ordinate descent rule as 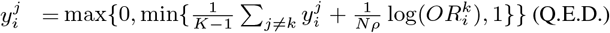.

The solutions can then be binarized by selecting an appropriate threshold like in the previous algorithm. An interesting observation is that the update rule for any *y^k^* is simply an average of all the other *y*’s and an additional term which is solely dependent on the odds ratio of the *k*th view. This can be implemented as seen in Algorithm 2:

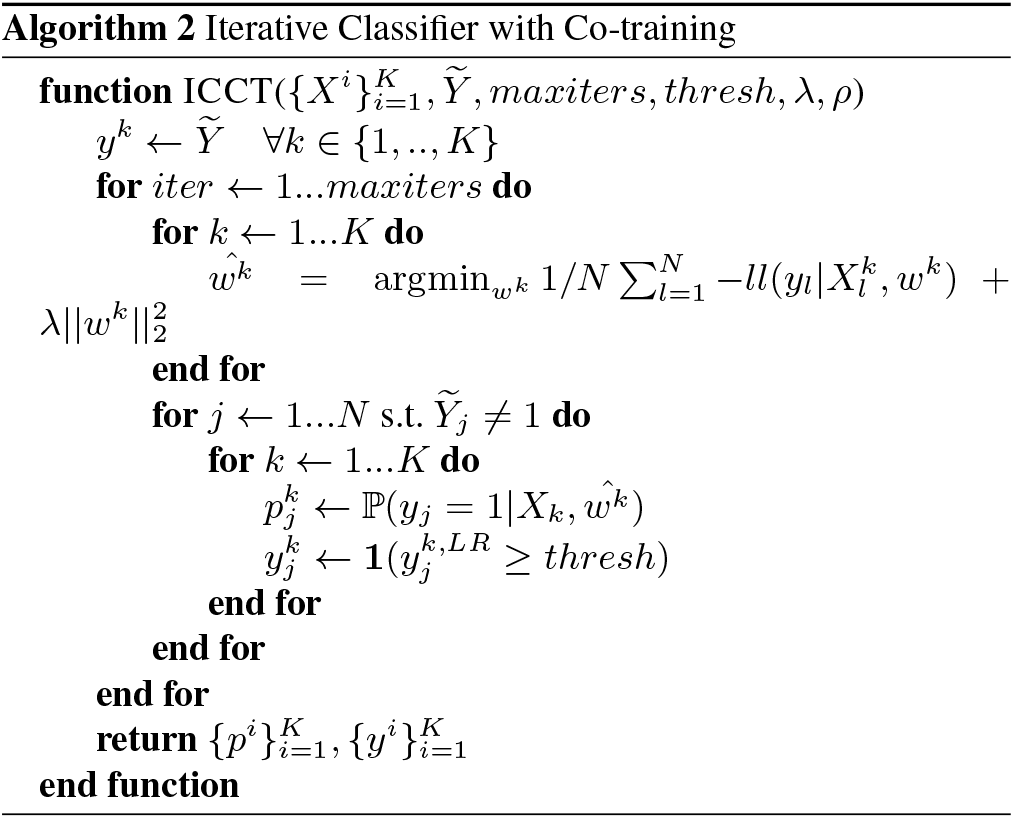

Similar to the separately trained approach, consensus is taken to obtain final multi-view predictions.

#### 2.3.3 Implementation and hyperparameter tuning

Both multiview iterative learning schemes were built using the Logistic regression in the *sci-kit learn* package of Python. A generalizable implementation of the code can be found at the link mentioned in the abstract. Values of *λ* for each feature set were chosen using a 10-fold crossvalidation approach using the original labels using the *LogisticRegressionCV* function in *sci-kit learn*. The value of *ρ* was chosen to be 1 /*N* for analysis of the RNA-Seq dataset based on performance on the benchmark datasets.

### 2.4 Evidence aggregated ranking

The goal of the evidence aggregated ranking scheme is to aggregate the predictions of the models trained using different featuresets and also (optionally) integrate unrelated external information from large sample GWAS studies. Here we develop a flexible scoring system that achieves the above stated goal:

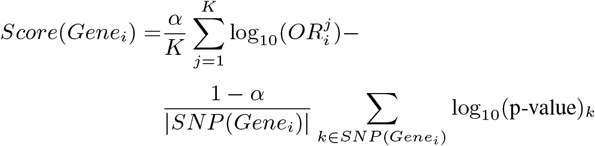

Here *α* ∈ (0, 1] is a user specified weighting parameter which controls the relative importance given to the external GWAS evidence vis-a-vis the model predictions using our featuresets. The models themselves are weighed equally relative to each other. For the purposes of this paper we chose the *α* = 0.5, thereby assigning equal weight to our model predictions and external GWAS evidence. The average of log transformed SNP p-value is chosen insteaf of the minimum p-value in order to capture the composite effect of all SNPs in a gene.

## 3 RESULTS

### 3.1 Comparison of learning approaches on benchmark datasets

To first test quantitatively test the relative efficiency of the two learning approaches, we first test them on some standard benchmark datasets obtained from *https://github.com/yeqinglee/mvdata* (used in (Li *et al*., 2015)):

**Handwritten digits**: This is a dataset containing handwritten digits (0 through 9) originally from UCI’s Machine Learning repository. It consists of 2000 data points. We use 3 of the published features namely: 240 pixel averages in 2 × 3 windows, 76 Fourier coefficients of the character shapes and 216 profile correlations.
**Caltech-101**: This is a dataset comprising of 7 classes of images amount to a total of 1474 images (Dueck and Frey (2007)). We use 3 of the published features namely: 48 Gabor features, 254 CENTRIST features and 40 features derived from Wavelet Momements.

For each dataset, we performed binary classification with different algorithms on each class separately, after corrupting the labels by randomly deleting 50% of the ‘true’ class labels to simulate the driver identification problem. The training was performed on corrupted labels while testing was performed on the actual labels. Algorithms were compared by their mean accuracy across all the class labels on the actual class labels. The algorithms compared were: i) Iterative classifiers trained on each feature type separately, ii) Iterative classifiers trained on each feature type separately followed by consensus among the learned models (using simple majority), and iii) Iterative classifier with co-training.

As seen in Figure 2, we see that Iterative classifier with co-training outperforms other algorithms on both standard datasets by a large margin, while Iterative classifier with consensus does not always lead to improvements over the best single view iteratively trained model. This is perhaps due to the difference in information content between the different views can sometimes make taking consensus ineffective.

**Fig. 2.**
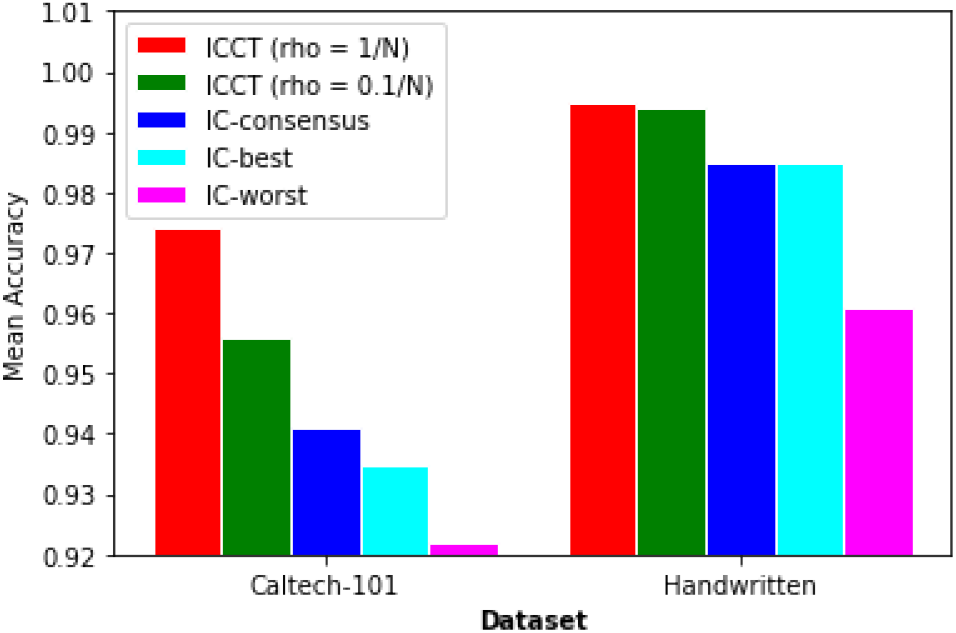
Comparison of various classification algorithms trained on corrupted class labels and tested on actual labels.

### 3.2 Validation of driver prediction using independent GWAS datasets

To validate our multi-view data aggregation schemes and generate a biologically meaningful ranking, we first generated gene-wise summary statistics from two seperate GWAS datasets, namely IGAP (Lambert *et al*., 2013) and Jansen (Jansen *et al*., 2019). The IGAP study has a sample size of 74,046 (25,580 cases and 48,466 controls) from individuals of European ancestry with over 7 million total SNPs. The Jansen study has a sample size of 455,258 (71,880 cases, 383,378 controls) also from European ancestry. This study contains in the addition to the data used in the IGAP study in addition to 3 complementary studies: Alzheimers Disease Sequencing Project (ADSP)), Psychiatric Genomics Consortium (PGC-ALZ) and UK Biobank studies.

For each of these GWAS datasets, we generated two gene-wise summary statistics, namely: i) mean of log p-value of SNPs (MLP) and ii) minimum p-value (MP) of SNPs. This was done by mapping each SNPs to a 10kb window around known protein coding gene locations in a reference gemome (hg38) and then computing the two summary statistics of interest per gene. The mapping of SNPs to genes was performed using the MAGMA software package (de Leeuw *et al*., 2015).

Similar to the benchmark datasets, we trained both IC and ICCT models on the three previously mentioned featuresets to obtain probabilities of all genes being driver genes for AD. In the absence of true labels for validation, we adopt a qualitative metric to test the model accuracies using external GWAS data. This was done by performing a Mann-Whitney U test between the distributions of MP values of predicted driver genes and genes not predicted to be drivers. A significant difference between the distributions would suggest that predicted driver genes contain more genes significant to AD than non-driver genes. Using this metric, we find that the ICCT-consensus model shows the strongest difference between the distributions (measure using the Mann-Whitney U p-value), followed by models trained on the network topological features trained as a part of the ICCT algorithm (Figure 3). It is seen in both datasets, that even some featureset specific predictions of the ICCT algorithm outperforms the basic iterative learning approach (IC), demonstrating the utility of co-training. Interestingly, the high relative performance of the network topological features when compared to the differential expression features implies that local and global network structure has a plays a strong role in determining which genes have causal effects on Alzheimers.

**Fig. 3.**
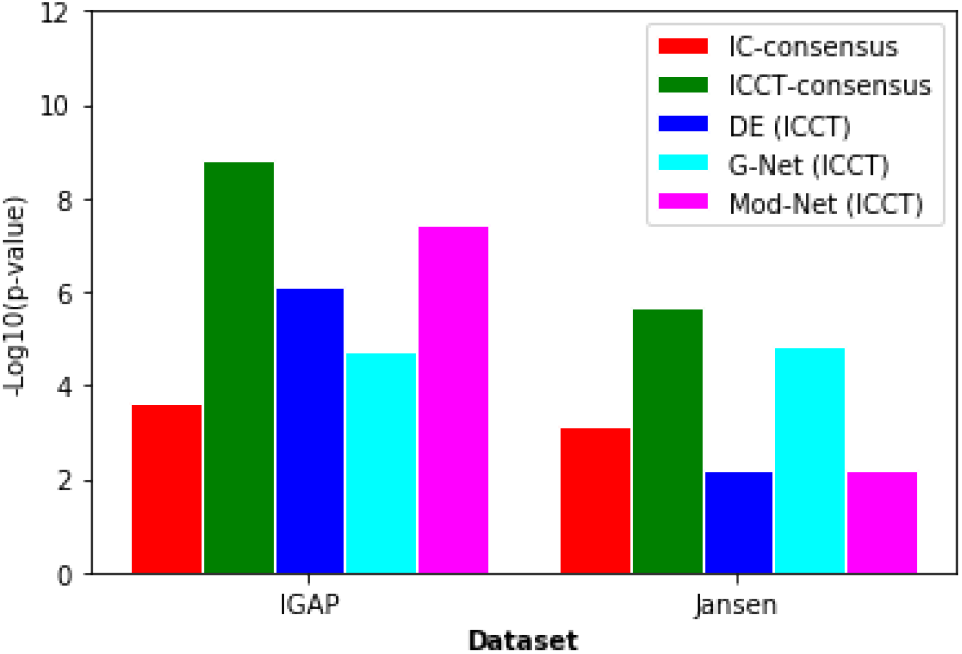
Results of the Mann-Whitney U test performed on IGAP and Jansen MP distributions for predicted driver vs non-driver genes.

### 3.3 Biological analysis of predicted drivers

Having demonstrated the statistical significance of the predicted driver genes, we ranked them using our ranking schema. The top 20 ranked genes can be seen in Table 2, which contains several genes strongly linked with AD such as APOE, APOC1, CD74, TREM2, SLC7A7 (Mukherjee *et al*., 2017; Ki *et al*., 2002; Kiyota *et al*., 2015; Jonsson *et al*., 2013) etc.. Table 2 also contains the minimum SNP p-values for each of these genes according to the IGAP and Jansen studies. It can be seen that while our models are not trained on any SNP information, the results strongly align with additional validation GWAS data.

**Table 2.** Top 20 ranked genes along with their associated driver score and minimum p-value from IGAP (Lambert *et al*., 2013) and Jansen et. al. (Jansen *et al*., 2019) GWAS datasets.

| Genes | Diver Score | Jansen p-value | IGAP p-value |
| --- | --- | --- | --- |
| APOC1 | 42.92 | <1E-308 | <1E-308 |
| APOE | 41.75 | <1E-308 | <1E-308 |
| BCAM | 5.88 | 1.60E-143 | 4.66E-69 |
| CD74 | 4.92 | 1.93E-02 | 1.20E-01 |
| TREM2 | 4.65 | 2.95E-15 | 1.07E-03 |
| CLPTM1 | 4.58 | 7.07E-50 | 2.80E-21 |
| DEF6 | 4.28 | 5.94E-03 | 3.52E-02 |
| SLC7A7 | 4.05 | 2.29E-03 | 2.36E-02 |
| DOCK2 | 3.72 | 9.14E-04 | 4.82E-03 |
| SPI1 | 3.62 | 1.06E-06 | 1.99E-06 |
| STEAP3 | 3.61 | 3.63E-05 | 2.21E-02 |
| PICALM | 3.56 | 2.19E-18 | 1.91E-12 |
| HMOX1 | 3.56 | 1.16E-02 | 1.43E-01 |
| CLU | 3.55 | 2.61E-19 | 2.48E-17 |
| MS4A6A | 3.55 | 1.55E-15 | 6.64E-11 |
| IRF5 | 3.45 | 1.21E-02 | 1.48E-02 |
| TYROBP | 3.44 | 1.34E-02 | 5.40E-02 |
| PARVG | 3.42 | 1.44E-02 | 1.05E-03 |
| ITGAL | 3.41 | 1.92E-04 | 4.36E-03 |
| PTPRC | 3.33 | 2.12E-03 | 7.24E-03 |

To further validate the results we performed gene set enrichment analyses with the top-500 ranked potential driver genes using Enrichr (Chen *et al*., 2013), a web based gene set enrichment tool. The top 20 significant processes and functions ranked according to their adjusted p-values can be seen in Table 3. Several of the processes such as immune response, amyloid processing, amyloid catablism, amyloid clearance, and apoptotic processes, and functions such as LDL binding and activity are already known to significantly altered in AD, whereas several other interesting ones such as endocytosis, scavenger receptor activity, and peptidase activity can lead to potential new insights into AD disease mechanisms.

**Table 3.**
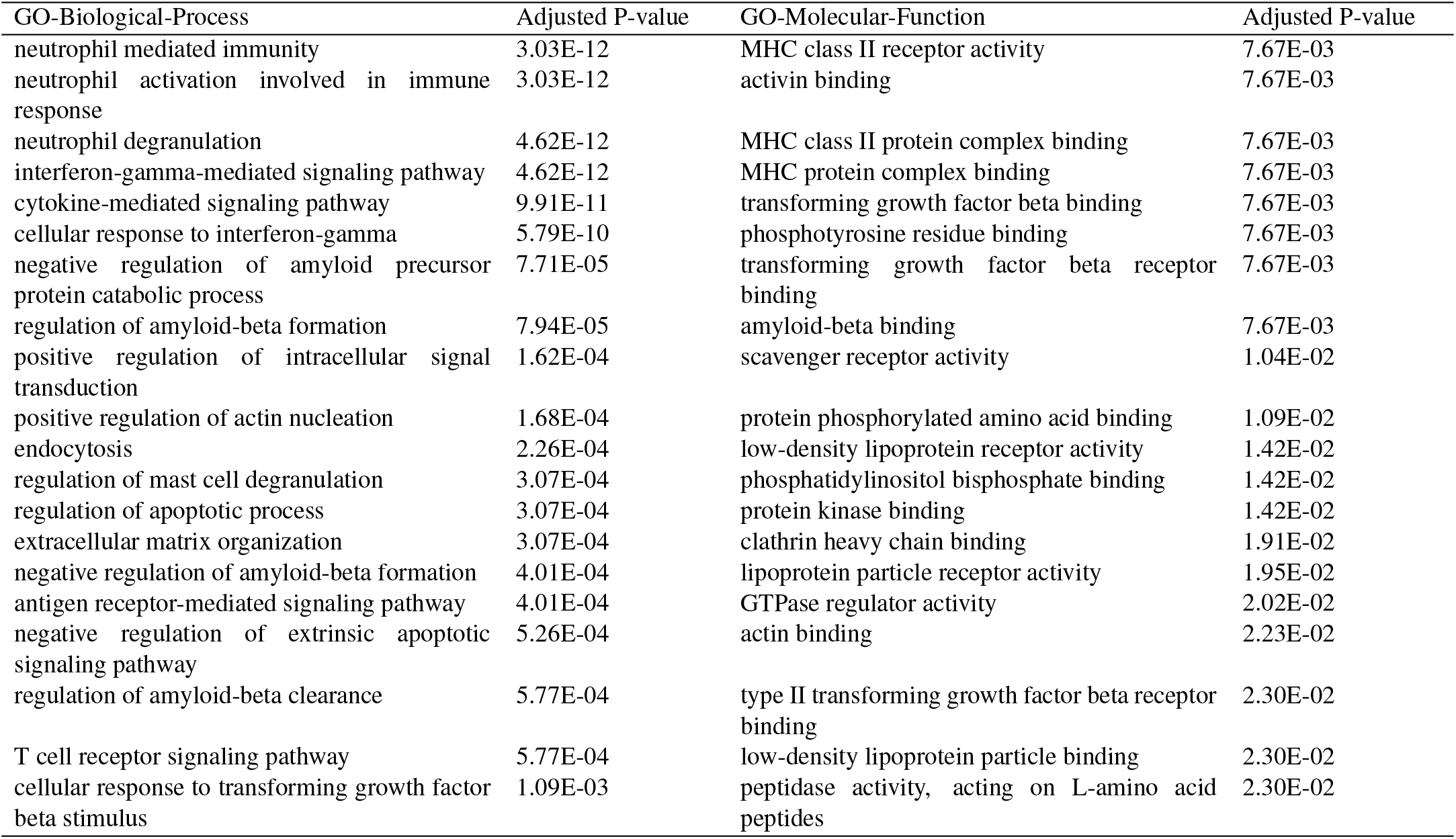
Top 20 enriched genesets for biological process and function along with their associated adjusted p-values obtained from Enrichr (Chen *et al*., 2013).

### 3.4 Analysis of top features for driver prediction models

Having noted that the network topological features provide are more predictive of the driver ranking of genes, we evaluate the most predictive features of each of the network featuresets in Table 4. We calculated the Spearman’s rank correlation for each feature with the model predictions for their featureset, to evaluate their relative predictive power. Interestingly, we find several highly correlated features from both featuresets. Upon closer look at the top 10 highly correlated features from the Module-Network featureset all are negatively correlated, with all the features derived from with DLPFC and TCX brain regions. This is intriguing because the sample size in DLPFC is largest (n=630), and the signal to noise ratio in TCX is highest (it is a highly affected brain region, and the median depth of sequencing for that study was 60 million reads compared to 35 million for the other studies). The same trend cannot be observed in the Global-Network featureset, where the top 10 features are associated with STG, PHG and DLPFC brain regions and all the correlations are positive. However, in this case, the top features are all associated with high connectivity of genes, which agrees with the popular notion that driver genes are also typically hub genes (Liu *et al*., 2012, 2011; Mukherjee *et al*., 2018). This can also be seen in Figure 4, where we note that most of the known drivers lie in one of the islands of genes (in the principle component plot) which corresponds to genes with very high degrees (or hubs).

**Fig. 4.**
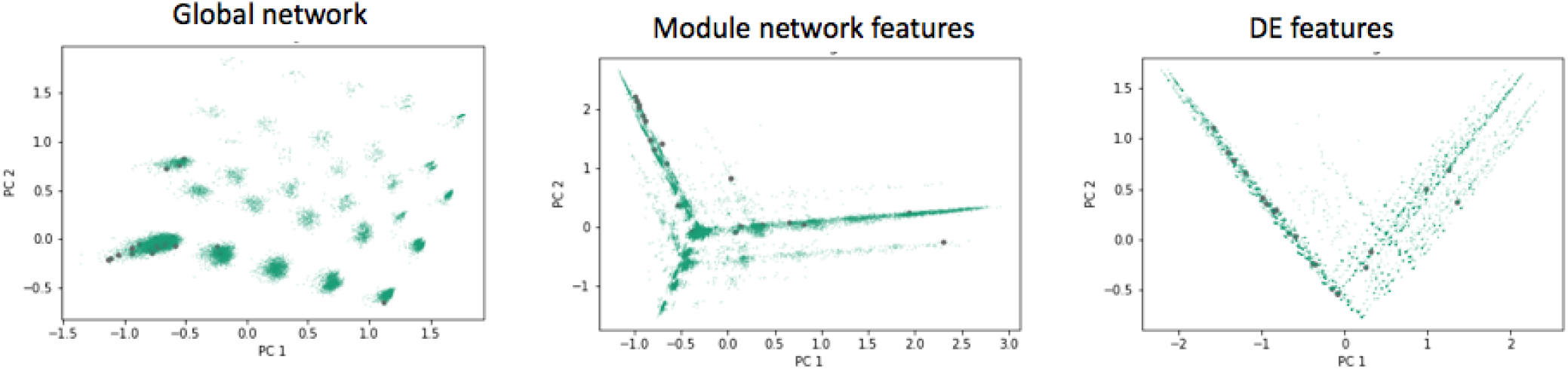
Known driver genes (colored in gray) and all other genes highlighted on the top two principal components for each of the three feature sets.

**Table 4.**
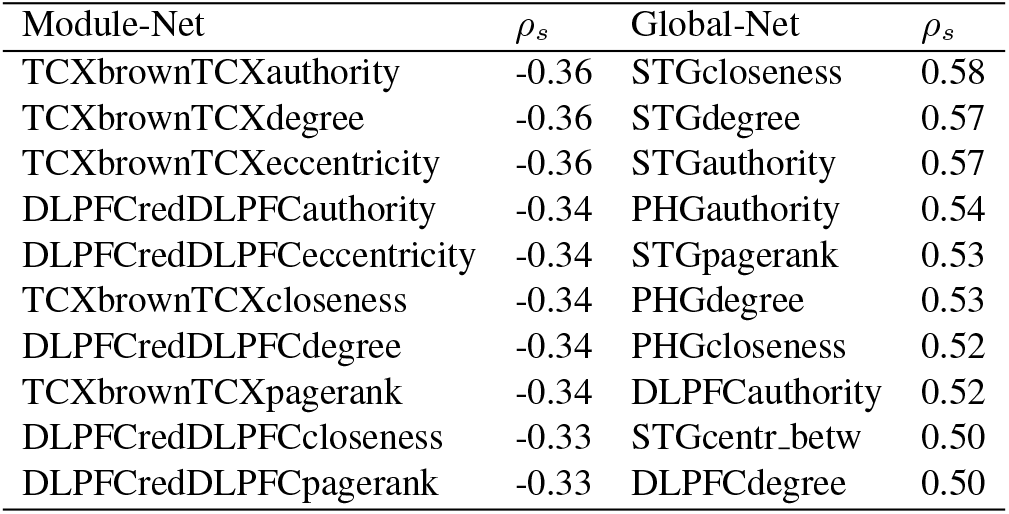
Spearman rank correlation (with model predictions) for the top 10 features of network topological feature sets.

## 4 CONCLUSION

Here, we provide a generalizable framework for integration of diverse systems biology outputs to rank and identify new transcriptomic and genetic drivers of Alzheimers disease. This provides evidence that integration of multiple systems biology resources can provide insights into new Alzheimers disease loci, which can help researchers prioritize future experimental studies focusing on specific genes and pathways that are driving disease etiology.

We currently demonstrate the utility of the approach on three RNA-Seq derived featuresets, providing strong qualitative agreement with known biology as well as previously published GWAS studies. Furthermore, we show the approach for driver gene prediction itself is a broadly application machine learning approach by demonstrating quantitative performance improvement over baseline models.

While the current work has focused on engineering and using RNA-Seq feature-sets, future work will focus on integrating other -omics datasets from the AMP-AD study to further improve the evidence driven ranking of driver genes. Another direction of future work will focus on identifying the relevance and agreement of different feature views. While the current approach equally weighs the predictions from different feature views, this may be unadvisable if a feature view has limited information about the driver-ness of genes.

